# Energy Guided Diffusion for Generating Neurally Exciting Images

**DOI:** 10.1101/2023.05.18.541176

**Authors:** Paweł A. Pierzchlewicz, Konstantin F. Willeke, Arne F. Nix, Pavithra Elumalai, Kelli Restivo, Tori Shinn, Cate Nealley, Gabrielle Rodriguez, Saumil Patel, Katrin Franke, Andreas S. Tolias, Fabian H. Sinz

## Abstract

In recent years, most exciting inputs (MEIs) synthesized from encoding models of neuronal activity have become an established method to study tuning properties of biological and artificial visual systems. However, as we move up the visual hierarchy, the complexity of neuronal computations increases. Consequently, it becomes more challenging to model neuronal activity, requiring more complex models. In this study, we introduce a new attention readout for a convolutional data-driven core for neurons in macaque V4 that outperforms the state-of-the-art task-driven ResNet model in predicting neuronal responses. However, as the predictive network becomes deeper and more complex, synthesizing MEIs via straightforward gradient ascent (GA) can struggle to produce qualitatively good results and overfit to idiosyncrasies of a more complex model, potentially decreasing the MEI’s model-to-brain transferability. To solve this problem, we propose a diffusion-based method for generating MEIs via Energy Guidance (EGG). We show that for models of macaque V4, EGG generates single neuron MEIs that generalize better across architectures than the state-of-the-art GA while preserving the within-architectures activation and requiring 4.7x less compute time. Furthermore, EGG diffusion can be used to generate other neurally exciting images, like most exciting natural images that are on par with a selection of highly activating natural images, or image reconstructions that generalize better across architectures. Finally, EGG is simple to implement, requires no retraining of the diffusion model, and can easily be generalized to provide other characterizations of the visual system, such as invariances. Thus EGG provides a general and flexible framework to study coding properties of the visual system in the context of natural images.^1^

## 1. Introduction

From the early works of Hubel and Wiesel [1], visual neuroscience has used the preferred stimuli of visual neurons to gain insight into the information processing in the brain. In recent years, deep learning has made big strides in predicting neuronal responses [2–15] enabling *in silico* stimulus synthesis of non-parametric most exciting inputs (MEIs) [16–18]. MEIs are images that strongly drive a selected neuron and can thus provide insights into its tuning properties. Up until now, they have been successfully used to find novel properties of neurons in various brain areas in mice and macaques [16–23].

However, as we move up the visual hierarchy, such as monkey visual area V4 and IT, the increasing non-linearity of neuronal responses with respect to the visual stimulus makes it more challenging to ❶ obtain models with high predictive performance for single neurons, and ❷ optimize perceptually plausible MEIs, that is, those not corrupted by adversarial high-frequency noise for example. Particularly, area V4 is known to be influenced by attention effects [24], and to be able to shift the location of receptive fields [25]. When models become more complex or units are taken from deeper layers of a network, existing MEI optimization methods based on gradient ascent (GA) can sometimes have difficulties producing qualitatively good results [26] and can overfit to the idiosyncrasies of more complex models, potentially decreasing the MEI’s model-to-brain transferability. Typically, these challenges are addressed by biasing MEIs towards the statistic of natural images, for instance by gradient pre-conditioning [26].

Here, we make two contributions towards the above points: ❶ We introduce a new attention readout for a convolutional data-driven core model for neurons in macaque V4 that outperforms the state-of-the-art robust task-driven ResNet [23, 27] model in predicting neuronal responses. ❷ To improve the quality of MEI synthesis we introduce a novel method for optimizing MEIs via Energy Guided Diffusion (EGG). EGG diffusion guides a pre-trained diffusion model with a learned neuronal encoding model to generate MEIs with a bias towards natural image statistics. Our proposed EGG method is simple to implement and, in contrast to similar approaches [28–30], requires no retraining of the diffusion model. We show that EGG diffusion not only yields MEIs that generalize better across architectures and are thus expected to drive real neurons equally well or better than GA-based MEIs but also provides a significant (4.7x) speed up over the standard GA method enhancing its utility for close-loop experiments such as inception loops [16, 17, 19, 23]. Since optimizing MEIs for thousands of neurons can take weeks [23], such a speed-up directly decreases the energy footprint of this technique. Moreover, we demonstrate that EGG diffusion straightforwardly generalizes to provide other characterizations of the visual system that can be phrased as an inverse problem, such as image reconstructions based on neuronal responses. The flexibility and generality of EGG thus make it a powerful tool for investigating the neural mechanisms underlying visual processing.

## 2 Attention readout for macaque area V4

### Background

Deep network based encoding models have set new standards in predicting neuronal responses to natural images [2–15]. Virtually all architectures of these encoding models consist of at least two parts: a *core* and a *readout*. The core is usually implemented via a convolutional network that extracts non-linear features Φ(***x***) from the visual input and is shared across all neurons to be predicted. It is usually trained through one of two paradigms: i) *task-driven*, where the core is pre-trained on a different task like object recognition [31, 32] and then only readout is trained to predict the neurons’ responses or ii) *data-driven* where the model is trained end-to-end to predict the neurons’ responses. The *readout* is a collection of predictors that map the core’s features to responses of individual neurons. With a few exceptions [33], the readout components and its parameters are neuron-specific and are therefore kept simple. Typically, the readout is implemented by a linear layer with a rectifying non-linearity. Different readouts differ by the constraints they put on the linear layer to reduce the number of parameters [3, 4, 33–35]. One key assumption all current readout designs make is that the readout mechanism does not change with the stimulus. In particular, this means that the location of the receptive field is fixed. While this assumption is reasonable for early visual areas like V1, it is not necessarily true for higher or mid-level areas such as macaque V4, which are known to be affected by attention effects and can even shift the location of the receptive fields [25]. This motivated us to create a more flexible readout mechanism for V4.

### State-of-the-art model: robust ResNet with Gaussian readout

In this study, we compare our data-driven model to a task-driven model [23], which is also composed of a *core* and *readout*. The core is a pre-trained robust ResNet50 [36, 37]. We use the layers up to layer 3 in the ResNet, providing a 1,024 dimension feature space. Then batch normalization is applied [38], followed by a ReLU non-linearity. The *Gaussian readout* [33] learns the position of each neuron and extracts a feature vector at this position. During training, the positions are sampled from a 2D Gaussian distribution with means *µ*_*n*_ and Σ_*n*_, during inference the *µ*_*n*_ positions are used. Then the extracted features are used in a linear non-linear model to predict neuronal responses.

### Proposed model: Data-driven core with attention readout

The predictive model is trained from scratch to predict the neuronal responses in an end-to-end fashion. Following Lurz et al. [33], the architecture is comprised of two main components. First, the *core*, a four-layer CNN with 64 channels per layer with an architecture identical to Lurz et al. [33]. Secondly, the attention *readout*, which builds upon the attention mechanism [39, 40] as it is used in the popular transformer architecture [41]. After adding a fixed positional embedding to [inedq] and normalization through LayerNorm [42] to get 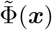 key and value embeddings are extracted from the core representation. This is done by position-wise linear projections 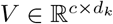 and 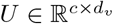 both of which have parameters shared across all neurons. Then, for each neuron a learned query vector 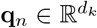 is compared with each position’s key embedding using scaled dot-product attention [41].

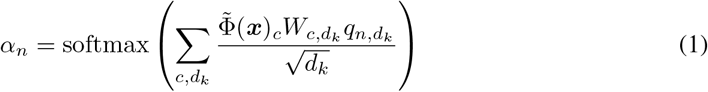

The result is a spatially normalized attention map *α*_*n*_ ∈ ℝ^*h×w×*1^ that indicates the most important feature locations for a neuron *n* for the input image. Using this attention map to compute a weighted sum of the value embeddings gives us a single feature vector for each neuron. A final neuron-specific affine projection with ELU non-linearity [43] gives rise to the predicted spike rate 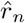(Fig. 2A). The model training is performed by minimizing the Poisson loss using the same setup as described in Willeke et al. [23].

**Figure 1:**
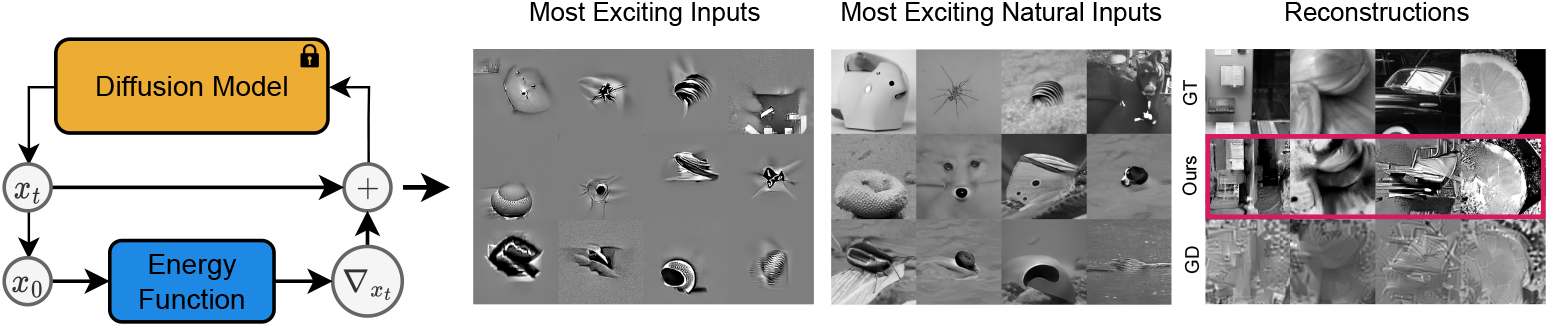
**Schematic** of the EGG diffusion method with a pre-trained diffusion model. Examples of applications: **Left**: Most Exciting Inputs for different neurons, **Middle:** Most Exciting Natural Inputs matched unit-wise to the MEIs. **Right**: Reconstructions in comparison to the ground truth (top) and gradient descent optimized (bottom).

**Figure 2:**
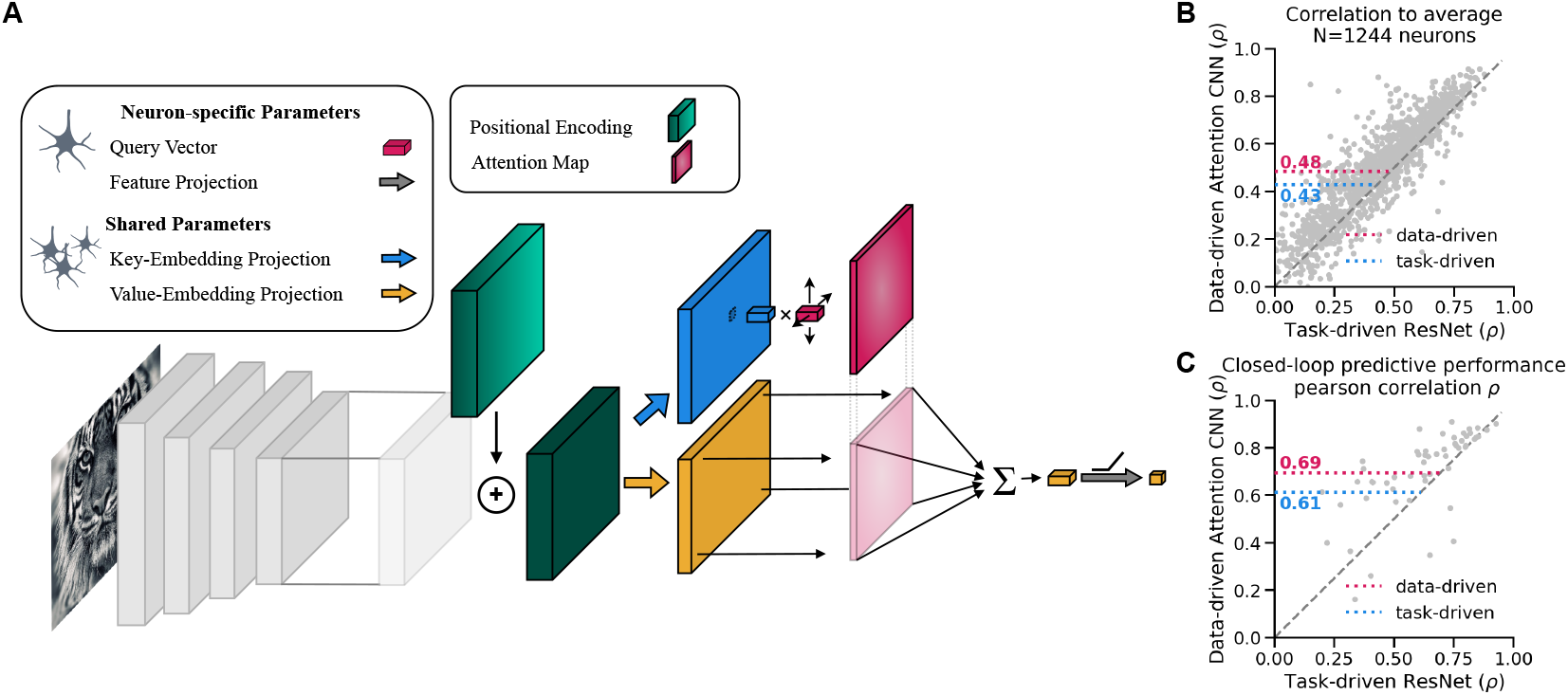
**a**) Schematic of the Attention Readout. **b**) Correlation to average scores for 1244 neurons. The data-driven with attention readout (pink) model shows a significant (as per the Wilcoxon signed rank test, p-value = 6.79 10^*−82*^) increase in the mean correlation to average in comparison to the task-driven ResNet (blue) model. **c**) Predictive performance comparison of the two models in a closed-loop MEI evaluation setting. Showing that the data-driven with attention readout model better predicts the in-vivo responses of the MEIs.

### Training data

We use the data from Willeke et al. [23] and briefly summarize their data acquisition in the supplementary materials section A.1.

### Results

Our attention readout on a CNN core significantly outperforms a state-of-the-art robust ResNet with a Gaussian readout in predicting neuronal responses of macaque V4 cells on unseen natural and model-derived images. We evaluate the model performance by the correlation between the model’s prediction and the averages of actual neuron responses across multiple presentations of a set of test images, as described by Willeke et al. [23]. We compared this predictive performance to a task-driven robust ResNet [37] on 1,244 individual neurons (Fig. 2B). The data-driven attention readout significantly outperforms the task-driven ResNet with Gaussian readout by 12% (Wilcoxon signed-rank test, p-value = 6.79 · 10^*−82*^). In addition, we evaluated the new readout on how well it predicts the real neuronal responses to 48 MEIs generated from the task-driven ResNet model [see 23] and 7 control natural images. Our data-driven CNN with the attention readout is better at predicting real neuronal responses, even for MEIs of another architecture (Fig. 2C). Please note that Willeke et al. [23] experimentally verified MEIs in only a subset of neurons and only used the neurons with high functional consistency across different experimental sessions. For that reason, we too can only compare the performance of model-derived MEIs on this subset of neurons.

## 3 Energy guided diffusion (EGG)

### 3.1 Algorithm and methods

In this section, we describe our approach to extract tuning properties of neuronal encoding models using a natural image prior as described by a diffusion model. In brief, we use previously established links between diffusion and score-based models and the fact that many tuning properties can be described as inverse problems (most exciting image, image reconstruction from neuronal activity, …) to combine an energy landscape defined by the neuronal encoding model with the energy landscape defined by the diffusion model and synthesize images via energy minimization. We show that this method leads to better generalization of MEIs and image reconstructions across architectures, faster generation, and allows for generating natural-looking stimuli.

#### Background: diffusion models

Recently, Denoising Diffusion Probabilistic Models (DDPMs) have proved to be successful at generating high-quality images [28, 44–49]. These models can be formalized as a variational autoencoder with a fixed encoder ***x***_0_ →***x***_*T*_ that turns a clean sample ***x***_0_ into a noisy one ***x***_*T*_ by repeated addition of Gaussian noise, and a learned decoder ***x***_*T*_→ ***x***_0_ [28], which is often described as inverting a diffusion process [44]. After training, the sampling process is initialized with a standard Normal sample ***x***_*T*_ ∼ 𝒩 (**0, I**) which is iteratively “denoised” for *T* steps until ***x***_0_ is reached. In the encoding, each step *t* corresponds to a particular noise level such that

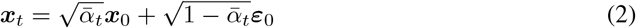

where 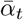 controls the signal strength at time *t* and *ε*_0_ ∼ 𝒩 (**0, I**) is independent Gaussian noise. In the decoding step, the diffusion model predicts the noise component *ε*_*θ*_(***x***_*t*_, *t*) at each step *t* of the diffusion process [28]. Then the sampling is performed according to

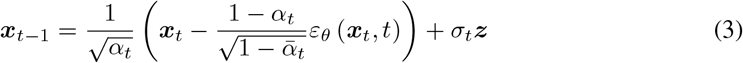

where ***z*** ∼ 𝒩 (**0, *I***).

Several previous works have established a link between diffusion models and energy-based models [50–52]. In particular, the diffusion model *ε*_*θ*_ (***x***_*t*_, *t*) can be interpreted as a *score function*, i.e. the gradient of a log-density or energy w.r.t. the data ∇_***x***_ log *p*(***x***) [53]. This link is particularly useful since combining two density models via a product is equivalent to adding their score functions.

#### EnerGy Guided Diffusion (EGG)

DDPMs learn to approximate the score of the distribution *p*(***x***_*t*_). The parameterization of diffusion models introduced by Ho et al. [28] only allows for the unconditioned generation of samples. Dhariwal and Nichol [46] introduced a method for sampling from a conditional distribution *p*_*t*_(***x***| ***y***), with diffusion models using a classifier *p*_*t*_(***y*** |***x***) known as classifier guidance. However, this method requires i) the classifier to be trained on the noisy images, and ii) is limited to conditions for which classification makes sense. At the core, this method relies on computing the score of the posterior distribution.

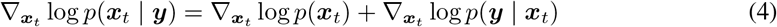

For classifier-guidance, the gradient of a model 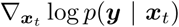 with respect to the noisy input ***x***_*t*_ is combined with the diffusion model 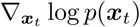 resulting in samples ***x***_0_ conditioned on the class ***y***. Note that this requires a model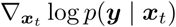 that has been trained on noisy samples of the diffusion before. Here we extend this approach to i) use neuronal encoding models, such as the ones described above, to guide the diffusion process and ii) to use a model trained on *clean* samples only. We achieve i) by defining conditioning as a sum of energies. Specifically, we redefine equation (4) in terms of the output of the diffusion model *ε*(***x***_*t*_, *t*) and an arbitrary energy function *E*(***x***_*t*_, *t*):

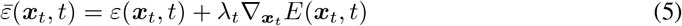

where *λ*_*t*_ is the energy scale. This takes advantage of the fact that sampling in DDPMs is functionally equivalent to Langevin dynamics Sohl-Dickstein et al. [44]. Langevin dynamics generally define the movement of particles in an energy field and in the special case when *E*(*x*) = −log *p*(*x*), Langevin dynamics generates samples from *p*(*x*). For this study, we use a constant value of *λ* and normalize the gradient of the energy function to a magnitude of 1.

To achieve ii) we use an approximate clean sample 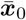 i.e. the original image, that can be estimated at each time step *t*. This is achieved by a simple trick, used in the code of Dhariwal and Nichol [46], of inverting the forward diffusion process, with the assumption that the predicted *ε*_*θ*_(***x***_*t*_, *t*) is the true noise:

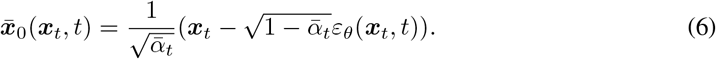

As a result, the energy function receives inputs that are in the domain of ***x***_0_ at much earlier time steps *t*, and hence makes it feasible to use energy functions only defined on ***x***_0_ and not ***x***_*t*_, dropping the requirement to provide an energy *E*(***x***_*t*_, *t*) that can take noisy images. This is particularly relevant in the domain of neural system identification, as encoding models are trained on neuronal responses to natural “clean” images [2–16, 20, 23, 33]. To get an energy that can understand noisy images would require showing the noisy images to the animals in experiments, which would make the use of this method prohibitively more difficult. Therefore, a guidance method that does not require training an additional model on noisy images allows researchers to apply EGG diffusion directly to existing models trained on neuronal responses and extract tuning properties from them.

#### Related work

Many other methods have been proposed to condition the samples of diffusion processes on additional information. Ho and Salimans [48] provided a method that addressed the second requirement of classifier-guidance by incorporating the condition ***y*** into the denoiser *ε*_*θ*_(***x***_*t*_, *t*, ***y***). However, to introduce a conditioning domain ***y*** in this classifier-free guidance, the whole diffusion model needs to be retrained. Furthermore, this link between diffusion models and energy-based models allowed several previous works to compose diffusion models to generate outputs that contain multiple desired aspects of a generated image [50–52]. However, these studies focus solely on generalizing the classifier-free guidance to allow guiding diffusion models with other diffusion models. While we were working on this project, Feng et al. [54] published a preprint where they used the score-based definition of diffusion models to introduce an image-based prior for inverse problems where the posterior score function is available. This work is most closely related to our approach. However, they focus on how to obtain samples and likelihoods from the true posterior. For that reason, they need guiding models to be proper score functions. We do not need that constraint and focus on guiding inverse problems defined by a more general energy function and focus particularly on the application to neuronal encoding models.

#### Image preprocessing for neural models

The neural models used in this study expect 100 ×100 images in grayscale. However, the output of the ImageNet pre-trained Ablated Diffusion Model (ADM) [46] is a 256 × 256 RGB image. We, therefore, use an additional compatibility step that performs i) downsampling from 256× 256 → 100× 100 with bilinear interpolation and ii) takes the mean across color channels providing the grayscale image. Each of these preprocessing steps is differentiable and is thus used end-to-end when generating the image.

### 3.2 Experiments

#### Most exciting images

We apply EGG diffusion to characterize properties of neurons in macaque area V4. For each of these experiments, we use the pre-trained ADM diffusion model trained on 256 ×256 ImageNet images from Dhariwal and Nichol [46]. In each of our experiments, we consider two paradigms: 1) **within** architecture, where we use two independently pre-trained ensembles containing 5 models of the same architecture (task-driven ResNet with Gaussian readout or data-driven with attention readout). We generate images on one and evaluate on the other. 2) **cross** architecture, two independently pre-trained ensembles containing 5 models of different architectures (ResNet and data-driven with attention readout). We demonstrate EGG on three tasks 1) Most Exciting Input (MEI) generation, where the generation method needs to generate an image that maximally excites an individual neuron, 2) Most Exciting Natural Image (MENI) generation, where a natural-looking image is generated that maximizes individual neuron responses, and 3) reconstruction of the input image from predicted neuronal responses. Running the experiments required a total of 7 GPU days. All computations were performed on a single consumer-grade GPU: NVIDIA GeForce RTX 3090 or NVIDIA GeForce RTX 2080 Ti depending on the availability.

MEIs have served as a powerful tool for visualizing features of a network, providing insights and testable predictions [16–20, 22, 55]. For the generation of MEIs, we selected 90 units at random from a subset of all 1,244 for which both ResNet and Attention CNN achieve at least a correlation of 0.5 to the average responses across repeated presentations. We compare our method to a vanilla gradient ascent (GA) method [23] which optimizes the pixels of an input image ***x*** to obtain the maximal response of the selected neuron. For the GA method, we use Gaussian blur preconditioning of the gradient. The stochastic gradient descent (SGD) optimizer was used with a learning rate of 10 and the image is optimized for 1,000 steps. We define EGG diffusion with the energy function 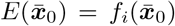 where *f*_*i*_ is the *i*-th neuron model and 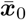 is the estimated clean sample. We optimize MEIs for both the ResNet model and the data-driven model with attention readout. We set the energy scale to *λ* = 10 for the ResNet and *λ* = 5 for the data-driven model with attention readout. *λ* was chosen via a grid search, for more details refer to Fig. 5B. The diffusion process was run for 100 respaced time steps for the task-driven model and 50 respaced time steps for the data-driven model. For both EGG and GA, we set the norm of the 100 ×100 image to a fixed value of 25. For each of the methods, we chose the best of 3 MEIs optimized from different seeds.

We show some examples of MEIs generated with EGG diffusion and GA for the two architectures in figure 3A. For more examples, refer to the supplementary materials figure S1. We find that the EGG-generated MEIs are significantly better (attention readout) or similarly (ResNet) activating within architectures and are significantly better at generalizing across architectures (Fig. 3B). This can also be observed by a significant increase in the mean activation across all units (Table 1). Perceptually, EGG-generated MEIs of the attention readout model looked more complex and natural than the GA-generated MEIs, and more similar to MEIs of the ResNet model trained on image classification.

**Table 1:** Comparison of the average unit activations in response to MEIs in two paradigms 1) within architectures and 2) cross architectures, for two architectures ResNet and Attention CNN. Bold marks the method which has higher mean activation, and the † marks the increases which are statistically significant (Wilcoxon signed-rank test, respective p-values: 0.08, 2.87·10^*−6*^, 2.84·10^*−10*^, 4.39 · 10^*−5*^).

| Paradigm | Within (ResNet) | Cross (ResNet) | Within (Attention) | Cross (Attention) |
| --- | --- | --- | --- | --- |
| Gradient Ascent | 11.43 | 5.51 | 7.59 | 4.42 |
| EGG Diffusion | **11.76** | **6.53<sup>†</sup>** | **10.56<sup>†</sup>** | **5.50<sup>†</sup>** |

**Figure 3:**
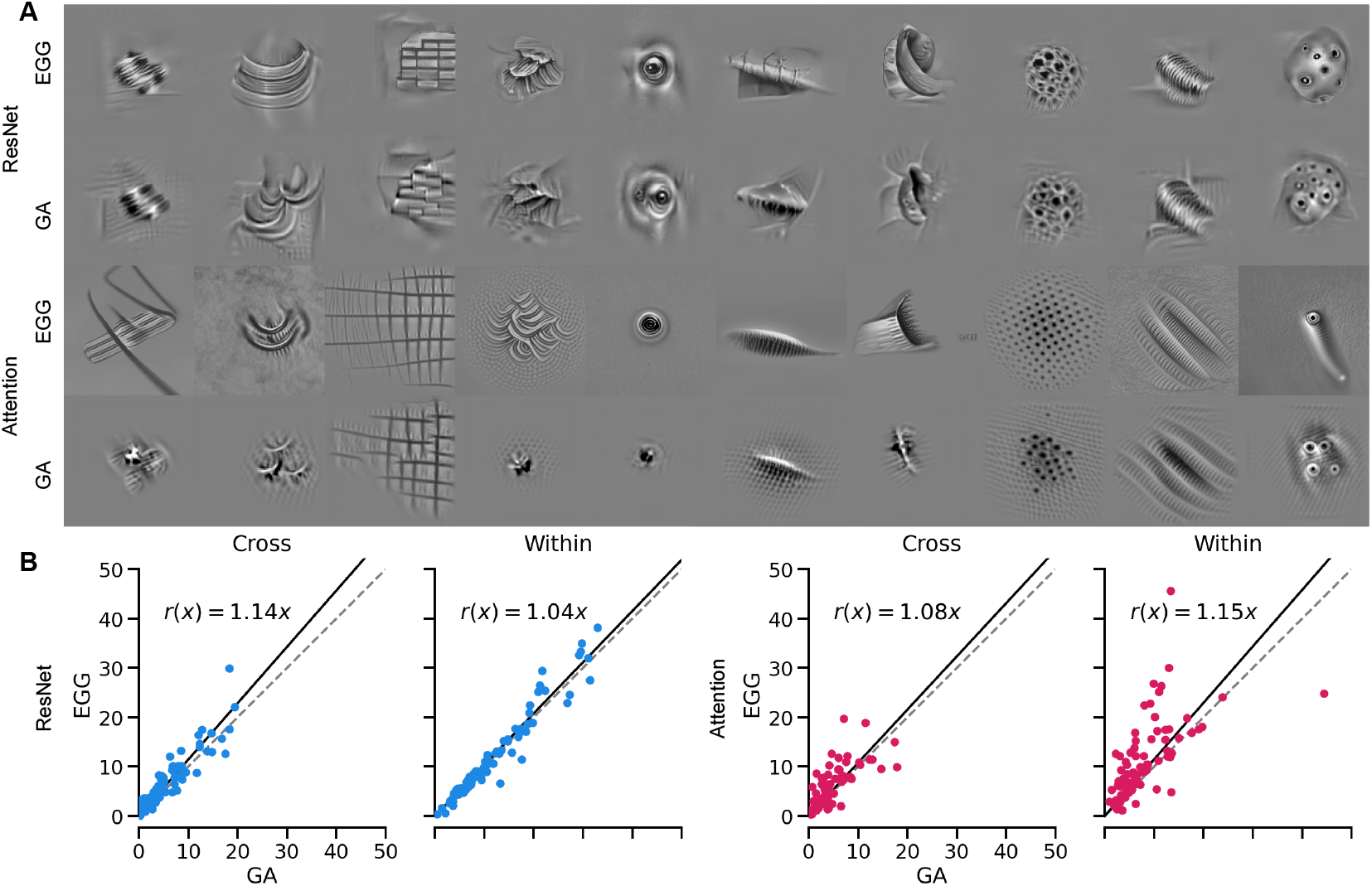
**a**) Examples of MEIs optimized using EGG diffusion and GA for macaque V4 ResNet and ACNN models. **b**) Comparison of activations for different neurons between EGG diffusion and GA on the Within and Cross Architecture validation paradigms. Line fits obtained via Huber regression with *ε* = 1.1.

Finally, EGG diffusion is almost 4.7-fold faster than GA, requiring only on average 46s per MEI in comparison to the required 219s for the GA method (Fig. 4) on a single NVIDIA GeForce RTX 3090 across 10 repetitions. This is a substantial gain, as Willeke et al. [23] required approximately 1.25 GPU years to optimize the MEIs presented in their study. With EGG, only approximately 0.25 GPU years would be needed to produce the results of the study, while providing higher quality and higher resolution MEIs. Thus, EGG can provide major savings in time and energy, *and* improve the quality of MEIs.

**Figure 4:**
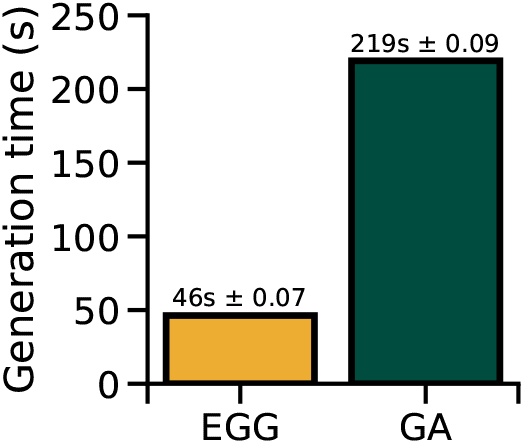
Mean comparison of the generation times between the EGG and GA (errorbars denote standard error).

**Figure 5:**
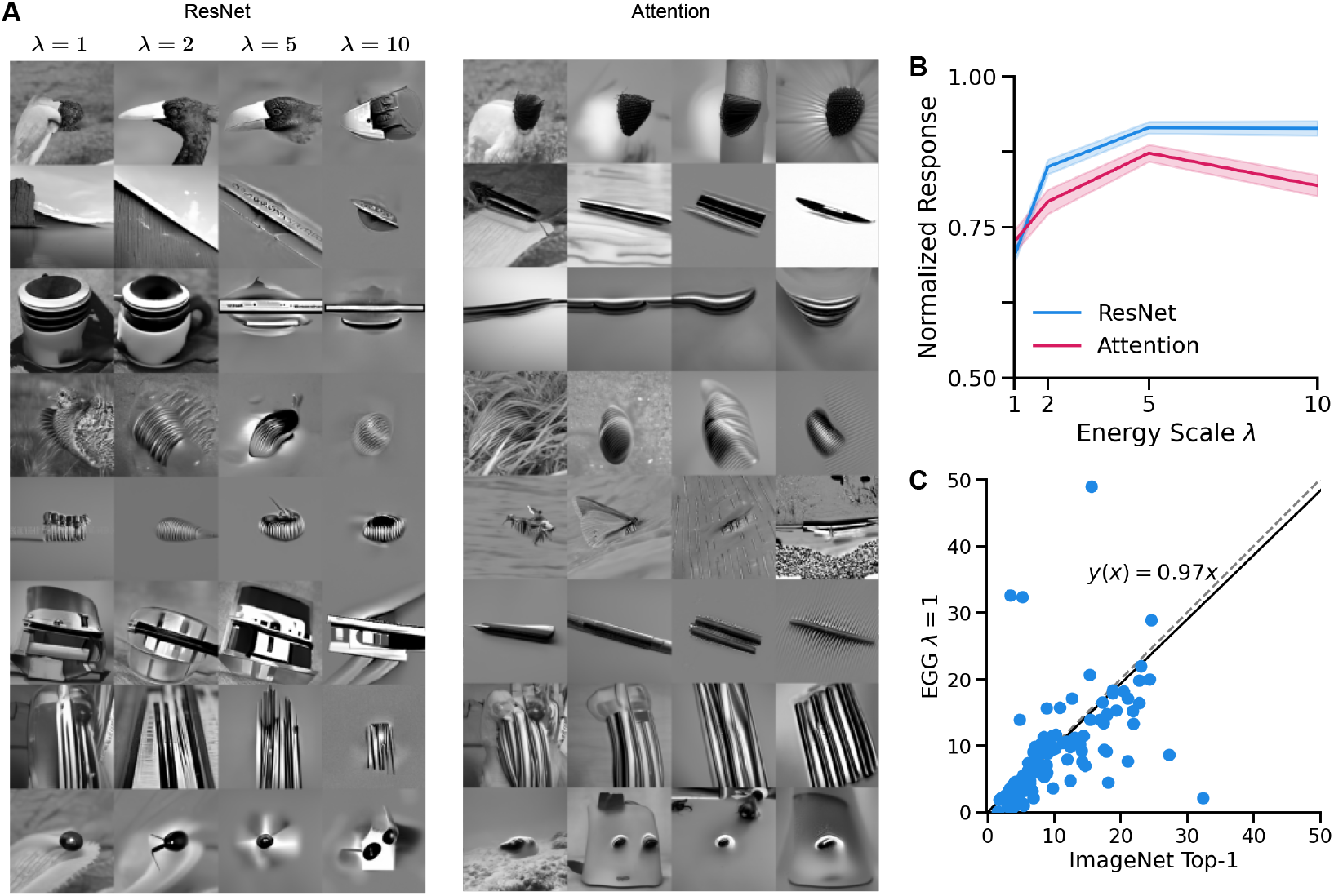
**a**) Examples of MENIs optimized using EGG diffusion in the macaque V4 for different neurons and different energy scales *λ* ∈{1, 2, 5, 10 }. **b**) Mean and standard error of the normalized activations of neurons across different energy scales. **c**) Comparison of the MENIs activations to the top-1 most activating ImageNet images per neuron in the cross-architecture domain. Line fit obtained via Huber regression with *ε* = 1.1. Three points at (11, 65), (9, 70), and (16, 120) are not shown for visualization purposes.

#### Controlling the “naturalness” to generate most exciting natural images

Unlike GA, EGG can also be used to synthesize more natural-looking stimuli by controlling the energy scale hyperparameter *λ*. Changing the value of *λ* trades off the importance of the maximization property of the image and its “naturalness”. To demonstrate this, we generated images for 150 neurons with the highest correlations to the average for the task-driven ResNet model with Gaussian readout. We used energy scales *λ*∈ {1, 2, 5, 10}, fixed the 100^2^ image norm to 50, and used 50 steps re-spaced from 1000. Each image was generated using 3 different seeds and the best-performing image on the generator model was selected.

We show examples of the generated images across different energy scales in figure 5A for both the task-driven ResNet with Gaussian readout and the data-driven with attention readout models. For more examples, refer to the supplementary material (Fig. S2, Fig. S3). Qualitatively, it can be observed that decreasing *λ* increases the naturalness of the generated image while preserving the features of the image that the neuron is tuned towards. We subsequently quantified the predicted responses across different values of *λ*. We find that increasing *λ* increases the predicted responses (Fig. 5B), however, at higher *λ* values the responses begin to plateau, or even decrease. Therefore, for generating MEIs with the ResNet model, we use *λ* = 10 and *λ* = 5 for the attention readout model.

Finally, we compare the generated MENIs (*λ* = 1) to a standard approach for finding natural images for individual neurons. To that end, we perform a search across 100k images from the ImageNet dataset [56] to find the top-1 most activating image for a particular unit. We then compare the predicted activations of the top-1 ImageNet image and the generated MENIs in the cross-architectures paradigm (Fig. 5C). We find that the generated MENIs drive comparable activation to the top-1 ImageNet images. Like in the MEI generation paradigm, EGG can thus significantly speed up the search for activating natural images, as it does not need to search through millions of images.

#### Image reconstruction from unit responses

Another application of EGG diffusion is image reconstruction from neuronal responses. A similar task has been attempted with success using diffusion models from human fMRI data [29, 30]. Given that only a small fraction of neurons were recorded, the image is encoded in an under-complete, significantly lower-dimensional space. Therefore, it is to be expected that the reconstructed image ***x*** will not necessarily be equal to the ground truth image ***x***_*gt*_. However, a better reconstruction ***x***^*∗*^ is one that generalizes across models. Therefore, regardless of the model *f* used, we should get ||*f* (***x***^*∗*^)− *f* (***x***_*gt*_)||_2_ = 0. This is trivially true for 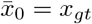 but, given the complexity of the model, there are likely other solutions.

We can reconstruct images in the EGG framework by defining the energy function as an *L*_2_ distance between the predicted responses to the ground truth image *f* (***x***_*gt*_) and the predicted responses to a generated image (Fig. 6A) *E*(***x***) = ||*f* (***x***) −*f* (***x***_*gt*_)||_2_. Note that, instead of *f* (***x***_*gt*_), we could also use recorded neuronal responses. The images are generated from the ResNet model with *λ* = 2 and 1000 timesteps, with the norm of the 100×100 image fixed to 60. We compare EGG to a GA method that simply minimizes the L2 distance. The GA uses an AdamW optimizer with a learning rate of 0.05. In GA, at each optimization step the image ***x***_*t*_ is Gaussian blurred and the norm is set to 60 before passing it to the neural encoding model. We optimize the GA reconstruction up to the point where the train *L*_2_ distance is matched between the GA and the EGG for a fair comparison of the generalization capabilities. We verified that the GA images do not improve qualitatively with more optimization steps (Fig. S5) We find that when generating the reconstruction using EGG diffusion we obtain 1) comparable within-architecture generalization and 2) much better cross-architecture generalization (Fig. 6B). The EGG-generated images produce lower within architecture distances for 84% of the images and for 98% in the cross-architecture case. We show examples of EGG diffusion and GA reconstructions in Fig. 6C. Qualitatively, the images optimized by EGG resemble the ground truth image much more faithfully than the GA images. More examples are available in the supplementary materials (Fig. S4).

**Figure 6:**
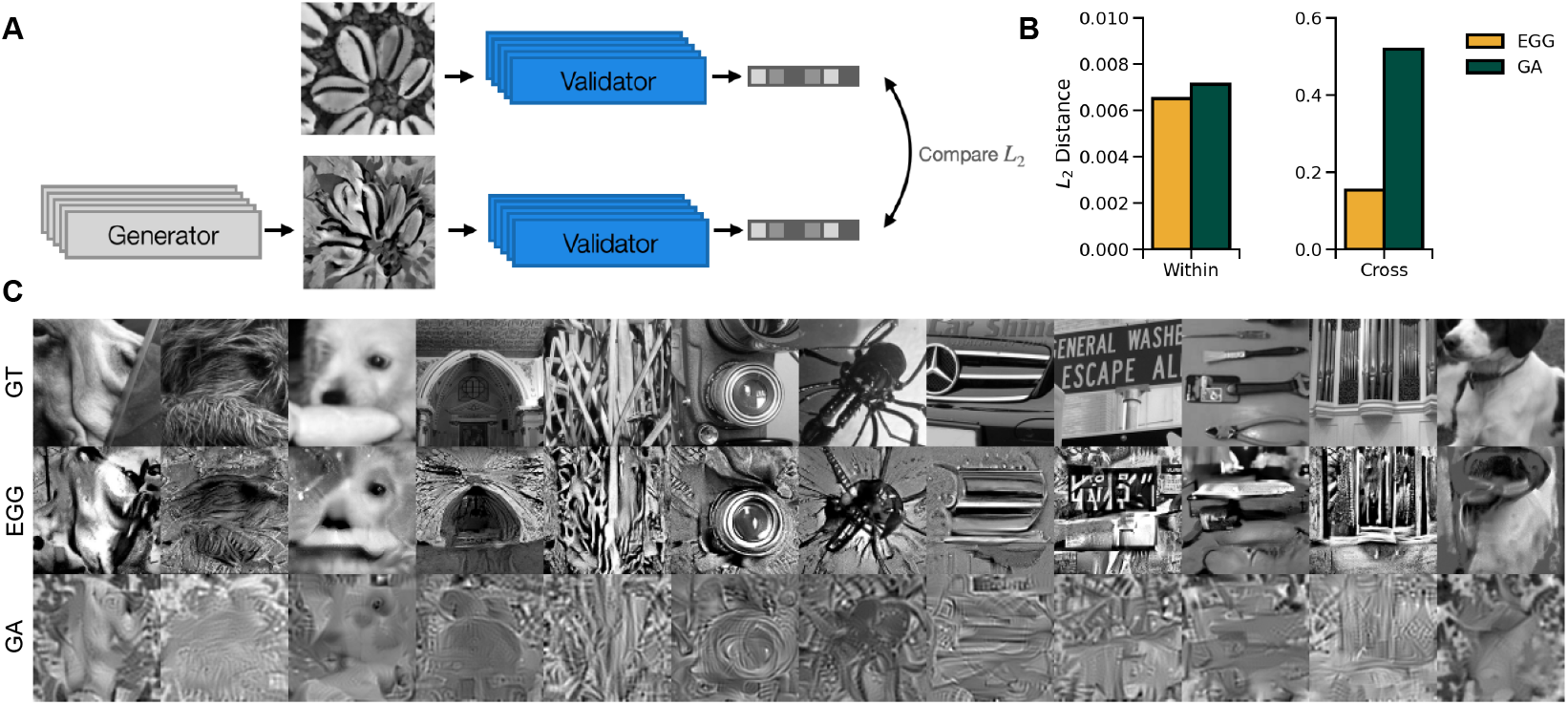
**a**) Schematic of the reconstruction paradigm. The generated image is compared to the ground truth image via *L*_2_ distance in the unit activations space. **b**) *L*_2_ distances in the unit activations space for the Within and Cross architecture domains comparing the EGG and GA generation methods. Shows that the EGG method generalizes better than GA across architectures. **c**) examples of reconstructions generated by EGG and GA in comparison to the ground truth (GT).

#### Limitations

While EGG diffusion on average performs better than GA it does come with limitations. Firstly, although the energy scale provides additional flexibility it is an additional hyperparameter that needs to be selected to obtain the desired results. Furthermore, the parameter value does not necessarily represent the same value same across energy functions. Secondly, the maximal number of steps to generate the sample is constricted by the pre-trained diffusion model. Finally, we identified that in 3 out of 90 cases, EGG diffusion failed to provide a satisfactory result with the ResNet model (Fig. 7), where GA was able to generate an MEI.

**Figure 7:**
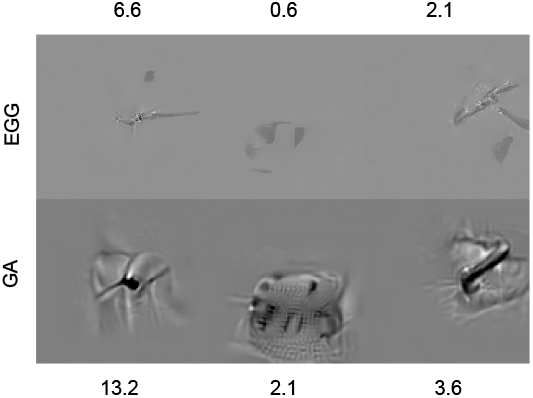
Examples of failure cases in comparison to the gradient ascent method. Text shows predicted response rate by the within-architecture validator.

## 4 Discussion

In this study, we introduced a new attention readout for a data-driven convolutional core model which provided the first data-driven model that outperformed task-driven models in predicting neurons in macaque V4. Furthermore, we propose a novel method for synthesizing images based on guiding diffusion models via energy functions (EGG). Our results indicate that EGG diffusion produces most exciting inputs (MEIs) which generalize better across architectures than the previous standard gradient ascent (GA) method. In addition, EGG diffusion significantly reduces compute time enabling larger-scale synthesis of visual stimuli. Comparing EGG-based MEIs to the ones found by Willeke et al. using GA, we find that the preferred image feature is usually preserved, but MEIs generated for the attention-readout model are more concentrated than their ResNet counterparts. This is not surprising, since the ResNet model was pre-trained on a large database of natural images. EGG diffusion is not limited to the generation of MEIs and, within the same framework, allows, among other characterizations, to 1) generate most exciting natural images (MENIs) which drive neurons on par with the most activating images in the ImageNet database, and 2) reconstruct images from unit responses, which generalize better across architectures and qualitatively resemble more the original image than images obtained via GA optimization. More generally, EGG can be used whenever the “constraint” on a particular image can be phrased in terms of an energy function. Thus EGG diffusion provides a flexible and powerful framework for studying coding properties of the visual system.

## Acknowledgments

The authors thank the International Max Planck Research School for Intelligent Systems (IMPRSIS) for supporting Konstantin Willeke and Arne Nix. The authors also thank Mohammad Bashiri and Suhas Shirinvasan for technical support and helpful discussions. The research was funded by the Carl-Zeiss-Stiftung (KW, FHS), the Cyber Valley Research Fund (AN, FHS). FHS is further supported by the German Federal Ministry of Education and Research (BMBF) via the Collaborative Research in Computational Neuroscience (CRCNS) (FKZ 01GQ2107), as well as the Collaborative Research Center (SFB 1233, Robust Vision) and the Cluster of Excellence “Machine Learning – New Perspectives for Science” (EXC 2064/1, project number 390727645). PP is supported by the German Federal Ministry for Economic Affairs and Climate Action (FKZ ZF4076506AW9). We also acknowledge support from the National Institute of Mental Health and National Institute of Neurological Disorders And Stroke under Award Number U19MH114830 and National Eye Institute award numbers R01 EY026927 and Core Grant for Vision Research T32-EY-002520-37 as well as the National Science Foundation Collaborative Research in Computational Neuroscience IIS-2113173.

## A Supplementary Material

### A.1 Training Data

Electrophysiological data were acquired as broadband signal (0.5Hz-16kHz), from a pair of male rhesus macaque monkeys (*Macaca mulatta*), using 32 channel linear silicon probes (NeuroNexus V1x32-Edge-10mm-60-177). The data was spike-sorted, and single units were isolated based on unit stability, refractory periods, and channel principal component pairs. Visual stimuli were presented to the animals on a 16:9 widescreen HD LCD monitor at 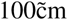 viewing distance. The animals were rewarded with juice if they maintained their gaze around a red fixation target throughout each trial. At the beginning of each recording session, the receptive fields (RFs) of the neurons were mapped in relation to a fixation target using sparse random dot stimuli, and the population RF was pulled towards the center of the screen by adjusting the fixation target. A collection of 24,075 images from ImageNet [56] was transformed into gray-scale and cropped to the central 420^2^ px and had 8 bit intensity resolution. These images were presented as visual stimuli during standalone generation recordings of 1244 units and during closed-loop recordings of 82 units. For details on the closed loop paradigm, please refer to Willeke et al. [23].

**Supplementary Figure S1:**
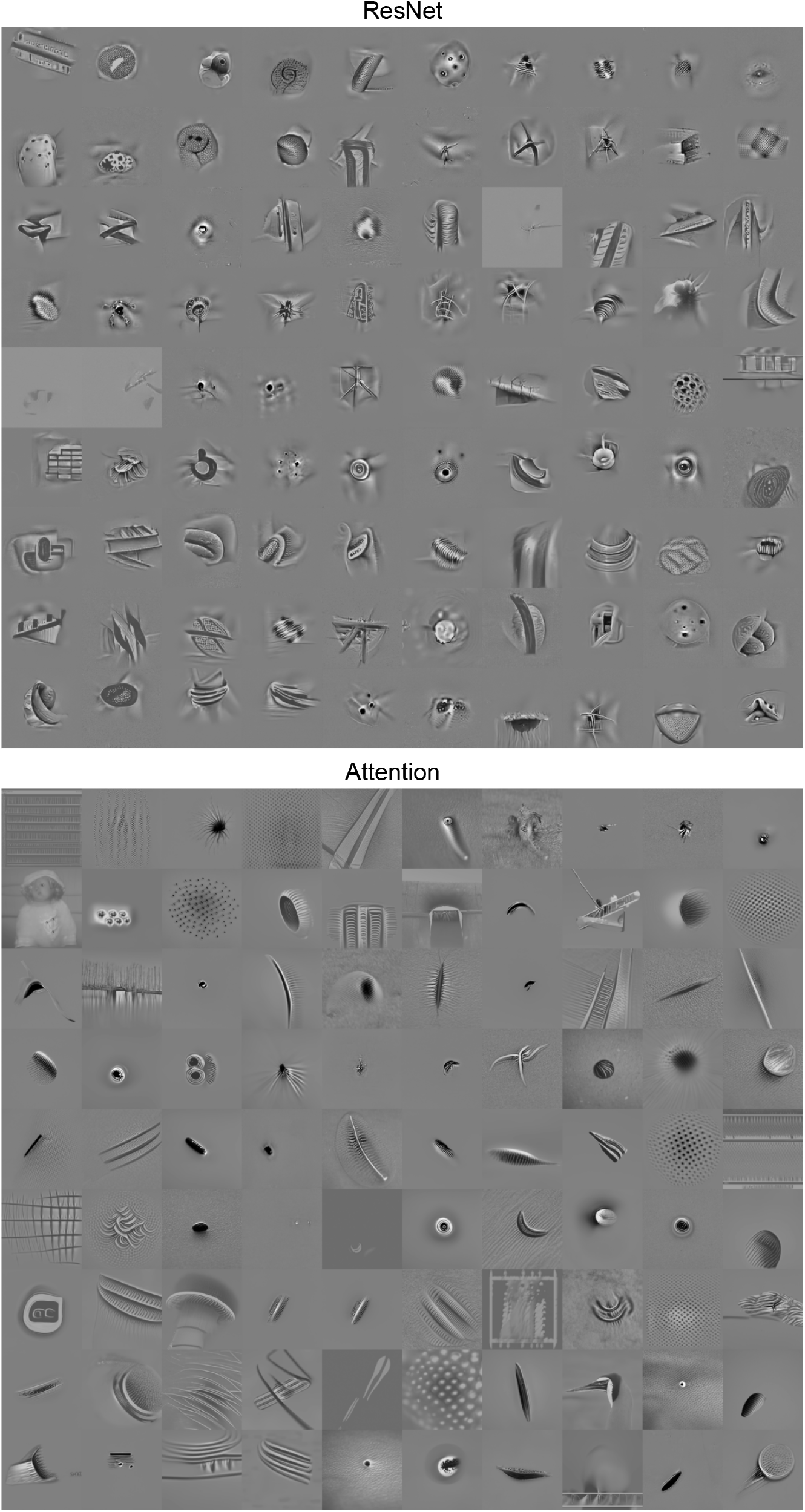
Examples of MEIs generated using EGG for the Attention readout and ResNet with Gaussian readout.

**Supplementary Figure S2:**
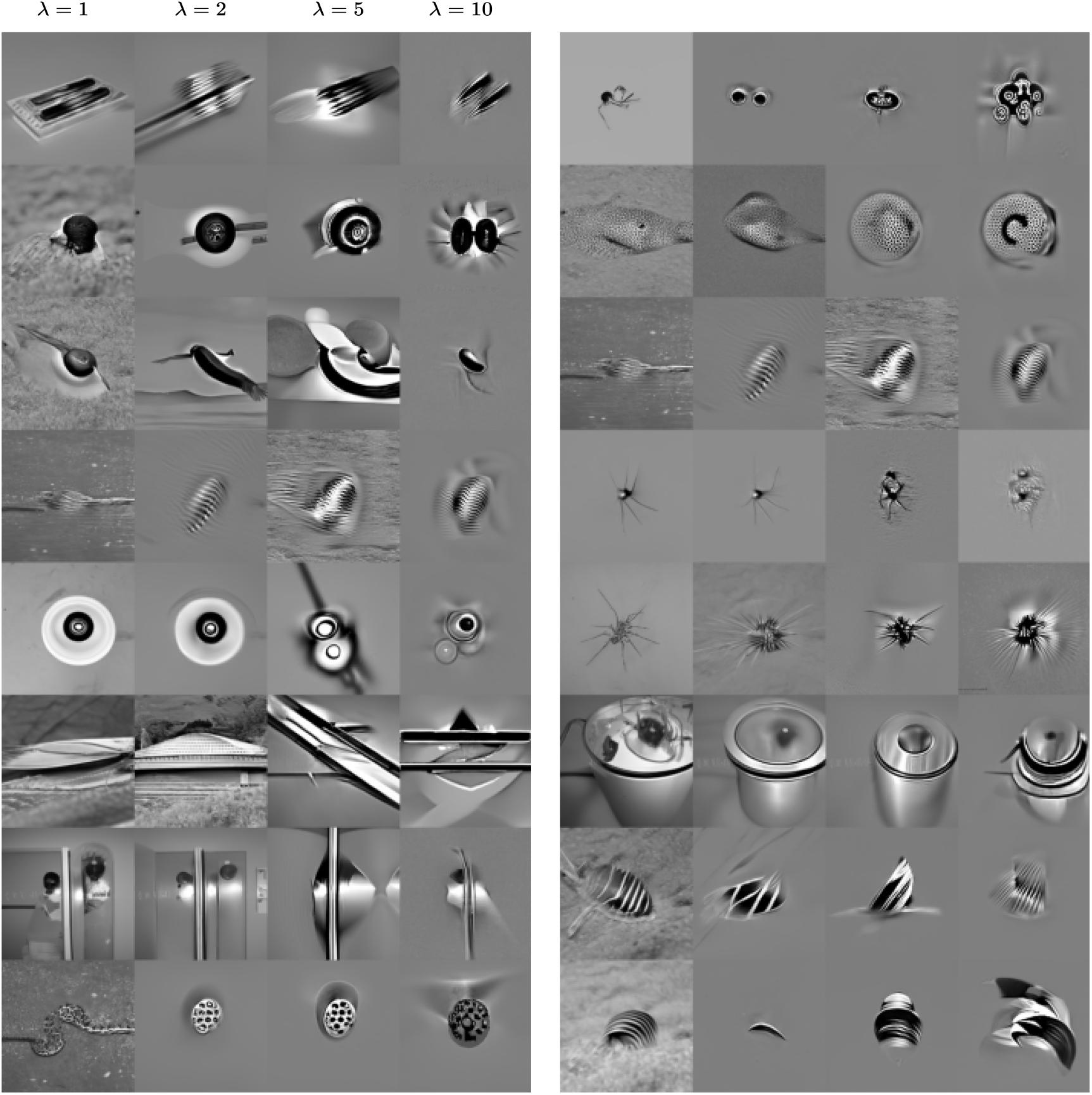
Examples of images generated using EGG diffusion in the Monkey V4 with different energy scales *λ* ∈{1, 2, 5, 10}. Generated for the ResNet model. Units not matched with the images shown for attention readout model.

**Supplementary Figure S3:**
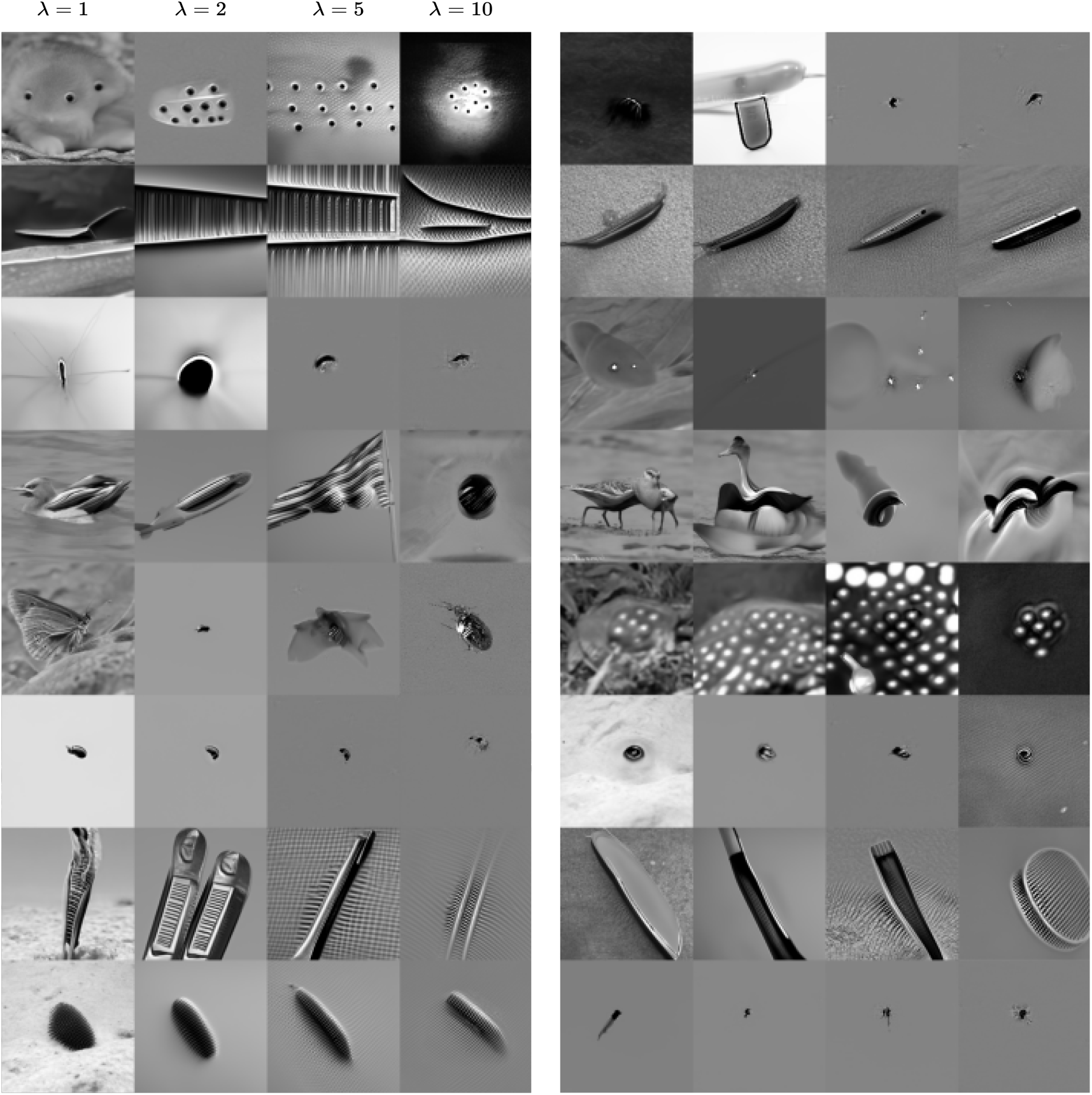
Examples of images generated using EGG diffusion in the Monkey V4 with different energy scales *λ* ∈{1, 2, 5, 10}. Generated for the data-drive with attention readout model. Units not matched with the images shown for ResNet model.

**Supplementary Figure S4:**
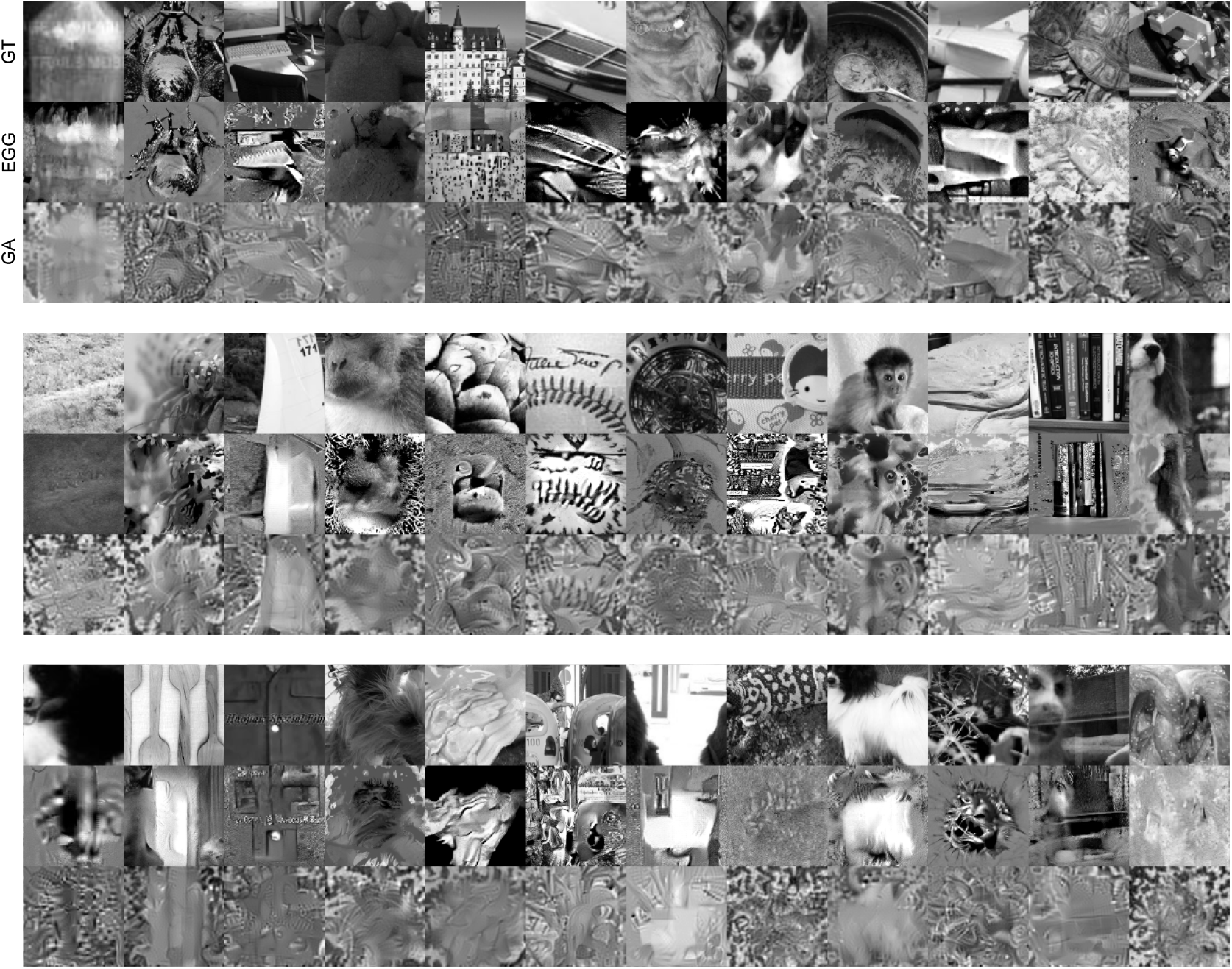
Reconstruction examples from the ResNet with Gaussian readout model. Generated using EGG diffusion and gradient descent.

**Supplementary Figure S5:**
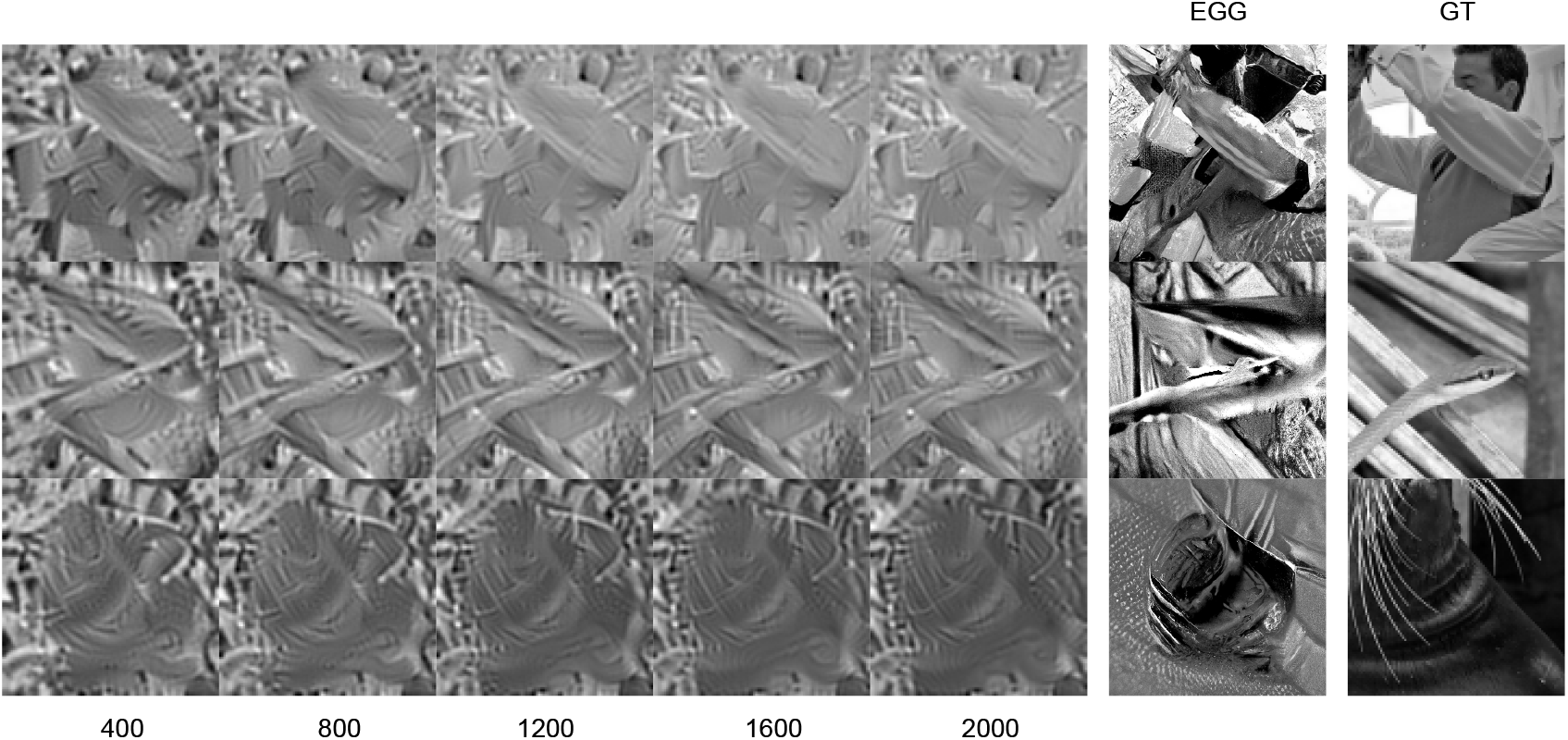
Examples of reconstructions using GD across various training lengths. Increasing the training does not bring the generated image closer visually to the GT nor EGG.

## Footnotes

1 The code is available at https://github.com/sinzlab/energy-guided-diffusion.

## References

[1] D H Hubel and T N Wiesel. Receptive fields of single neurones in the cat’s striate cortex. J. Physiol., 148(3):574–591, October 1959.

[2] Charles F Cadieu, Ha Hong, Daniel L K Yamins, Nicolas Pinto, Diego Ardila, Ethan A Solomon, Najib J Majaj, and James J DiCarlo. Deep neural networks rival the representation of primate IT cortex for core visual object recognition. PLoS Comput. Biol., 10(12):e1003963, December 2014.

[3] Daniel L K Yamins, Ha Hong, Charles F Cadieu, Ethan A Solomon, Darren Seibert, and James J DiCarlo. Performance-optimized hierarchical models predict neural responses in higher visual cortex. Proc. Natl. Acad. Sci. U. S. A., 111(23):8619–8624, June 2014.

[4] Santiago A Cadena, George H Denfield, Edgar Y Walker, Leon A Gatys, Andreas S Tolias, Matthias Bethge, and Alexander S Ecker. Deep convolutional models improve predictions of macaque V1 responses to natural images. PLoS Comput. Biol., 15(4):e1006897, April 2019.

[5] Fabian H Sinz, Alexander S Ecker, Paul G Fahey, Edgar Y Walker, Erick Cobos, Emmanouil Froudarakis, Dimitri Yatsenko, Xaq Pitkow, Jacob Reimer, and Andreas S Tolias. Stimulus domain transfer in recurrent models for large scale cortical population prediction on video. In Proceedings of the 32nd International Conference on Neural Information Processing Systems, NIPS’18, pages 7199–7210, Red Hook, NY, USA, December 2018. Curran Associates Inc.

[6] Yimeng Zhang, Tai Sing Lee, Ming Li, Fang Liu, and Shiming Tang. Convolutional neural network models of V1 responses to complex patterns. J. Comput. Neurosci., 46(1):33–54, February 2019.

[7] Lane T McIntosh, Niru Maheswaranathan, Aran Nayebi, Surya Ganguli, and Stephen A Baccus. Deep learning models of the retinal response to natural scenes. In Proceedings of the 30th International Conference on Neural Information Processing Systems, NIPS’16, pages 1369–1377, Red Hook, NY, USA, December 2016. Curran Associates Inc.

[8] David A Klindt, Alexander S Ecker, Thomas Euler, and Matthias Bethge. Neural system identification for large populations separating “what” and “where”. November 2017.

[9] William F Kindel, Elijah D Christensen, and Joel Zylberberg. Using deep learning to probe the neural code for images in primary visual cortex. J. Vis., 19(4):29, April 2019.

[10] Alexander S Ecker, Fabian H Sinz, Emmanouil Froudarakis, Paul G Fahey, Santiago A Cadena, Edgar Y Walker, Erick Cobos, Jacob Reimer, Andreas S Tolias, and Matthias Bethge. A rotation-equivariant convolutional neural network model of primary visual cortex. September 2018.

[11] Benjamin R Cowley and Jonathan W Pillow. High-contrast “gaudy” images improve the training of deep neural network models of visual cortex. June 2020.

[12] Max F Burg, Santiago A Cadena, George H Denfield, Edgar Y Walker, Andreas S Tolias, Matthias Bethge, and Alexander S Ecker. Learning divisive normalization in primary visual cortex. PLoS Comput. Biol., 17(6):e1009028, June 2021.

[13] Eleanor Batty, Josh Merel, Nora Brackbill, Alexander Heitman, Alexander Sher, Alan Litke, E J Chichilnisky, and Liam Paninski. Multilayer recurrent network models of primate retinal ganglion cell responses. July 2022.

[14] Mohammad Bashiri, Edgar Walker, Konstantin-Klemens Lurz, Akshay Jagadish, Taliah Muhammad, Zhiwei Ding, Zhuokun Ding, Andreas Tolias, and Fabian Sinz. A flow-based latent state generative model of neural population responses to natural images. Adv. Neural Inf. Process. Syst., 34:15801–15815, December 2021.

[15] Ján Antolík, Sonja B Hofer, James A Bednar, and Thomas D Mrsic-Flogel. Model constrained by visual hierarchy improves prediction of neural responses to natural scenes. PLoS Comput. Biol., 12(6):e1004927, June 2016.

[16] Edgar Y Walker, Fabian H Sinz, Erick Cobos, Taliah Muhammad, Emmanouil Froudarakis, Paul G Fahey, Alexander S Ecker, Jacob Reimer, Xaq Pitkow, and Andreas S Tolias. Inception loops discover what excites neurons most using deep predictive models. Nat. Neurosci., 22(12): 2060–2065, December 2019.

[17] Pouya Bashivan, Kohitij Kar, and James J DiCarlo. Neural population control via deep image synthesis. Science, 364(6439), 2019.

[18] Carlos R Ponce, Will Xiao, Peter F Schade, Till S Hartmann, Gabriel Kreiman, and Margaret S Livingstone. Evolving images for visual neurons using a deep generative network reveals coding principles and neuronal preferences. Cell, 177(4):999–1009.e10, 2019.

[19] Katrin Franke, Konstantin F Willeke, Kayla Ponder, Mario Galdamez, Na Zhou, Taliah Muhammad, Saumil Patel, Emmanouil Froudarakis, Jacob Reimer, Fabian H Sinz, and Andreas S Tolias. State-dependent pupil dilation rapidly shifts visual feature selectivity. Nature, 610(7930): 128–134, October 2022.

[20] Larissa Höfling, Klaudia P Szatko, Christian Behrens, Yongrong Qiu, David A Klindt, Zachary Jessen, Gregory W Schwartz, Matthias Bethge, Philipp Berens, Katrin Franke, Alexander S Ecker, and Thomas Euler. A chromatic feature detector in the retina signals visual context changes. December 2022.

[21] Jiakun Fu, Suhas Shrinivasan, Kayla Ponder, Taliah Muhammad, Zhuokun Ding, Eric Wang, Zhiwei Ding, Dat T Tran, Paul G Fahey, Stelios Papadopoulos, Saumil Patel, Jacob Reimer, Alexander S Ecker, Xaq Pitkow, Ralf M Haefner, Fabian H Sinz, Katrin Franke, and Andreas S Tolias. Pattern completion and disruption characterize contextual modulation in mouse visual cortex. March 2023.

[22] Zhiwei Ding, Dat T Tran, Kayla Ponder, Erick Cobos, Zhuokun Ding, Paul G Fahey, Eric Wang, Taliah Muhammad, Jiakun Fu, Santiago A Cadena, Stelios Papadopoulos, Saumil Patel, Katrin Franke, Jacob Reimer, Fabian H Sinz, Alexander S Ecker, Xaq Pitkow, and Andreas S Tolias. Bipartite invariance in mouse primary visual cortex. March 2023.

[23] Konstantin F Willeke, Kelli Restivo, Katrin Franke, Arne F Nix, Santiago A Cadena, Tori Shinn, Cate Nealley, Gabby Rodriguez, Saumil Patel, Alexander S Ecker, Fabian H Sinz, and Andreas S Tolias. Deep learning-driven characterization of single cell tuning in primate visual area V4 unveils topological organization. May 2023.

[24] Tirin Moore, Katherine M Armstrong, and Mazyar Fallah. Visuomotor origins of covert spatial attention. Neuron, 40(4):671–683, November 2003.

[25] A S Tolias, T Moore, S M Smirnakis, E J Tehovnik, A G Siapas, and P H Schiller. Eye movements modulate visual receptive fields of V4 neurons. Neuron, 29(3):757–767, March 2001.

[26] Chris Olah, Alexander Mordvintsev, and Ludwig Schubert. Feature visualization. Distill, 2(11): e7, November 2017.

[27] Yifei Ren and Pouya Bashivan. How well do models of visual cortex generalize to out of distribution samples? May 2023.

[28] Jonathan Ho, Ajay Jain, and Pieter Abbeel. Denoising diffusion probabilistic models. Adv. Neural Inf. Process. Syst., 33:6840–6851, 2020.

[29] Yu Takagi and Shinji Nishimoto. High-resolution image reconstruction with latent diffusion models from human brain activity. March 2023.

[30] Yizhuo Lu, Changde Du, Dianpeng Wang, and Huiguang He. MindDiffuser: Controlled image reconstruction from human brain activity with semantic and structural diffusion. March 2023.

[31] Santiago A Cadena, Konstantin F Willeke, Kelli Restivo, George Denfield, Fabian H Sinz, Matthias Bethge, Andreas S Tolias, and Alexander S Ecker. Diverse task-driven modeling of macaque V4 reveals functional specialization towards semantic tasks. May 2022.

[32] Daniel L K Yamins and James J DiCarlo. Using goal-driven deep learning models to understand sensory cortex. Nat. Neurosci., 19(3):356–365, March 2016.

[33] Konstantin-Klemens Lurz, Mohammad Bashiri, Konstantin Willeke, Akshay K Jagadish, Eric Wang, Edgar Y Walker, Santiago A Cadena, Taliah Muhammad, Erick Cobos, Andreas S Tolias, Alexander S Ecker, and Fabian H Sinz. Generalization in data-driven models of primary visual cortex. April 2021.

[34] D A Klindt, A S Ecker, T Euler, and M Bethge. Neural system identification for large populations separating “what” and “where”. In Advances in Neural Information Processing Systems, pages 4–6, 2017.

[35] F Sinz, A S Ecker, P Fahey, E Walker, E Cobos, E Froudarakis, D Yatsenko, X Pitkow, J Reimer, and A Tolias. Stimulus domain transfer in recurrent models for large scale cortical population prediction on video. In Advances in Neural Information Processing Systems 31. 2018.

[36] Kaiming He, Xiangyu Zhang, Shaoqing Ren, and Jian Sun. Deep residual learning for image recognition. December 2015.

[37] Hadi Salman, Andrew Ilyas, Logan Engstrom, Ashish Kapoor, and Aleksander Madry. Do adversarially robust ImageNet models transfer better? July 2020.

[38] Sergey Ioffe and Christian Szegedy. Batch normalization: Accelerating deep network training by reducing internal covariate shift. February 2015.

[39] Dzmitry Bahdanau, Kyunghyun Cho, and Yoshua Bengio. Neural machine translation by jointly learning to align and translate. arXiv preprint 1409.0473, 2014.

[40] Alex Graves. Generating sequences with recurrent neural networks. arXiv preprint 1308.0850, 2013.

[41] Ashish Vaswani, Noam Shazeer, Niki Parmar, Jakob Uszkoreit, Llion Jones, Aidan N Gomez, Łukasz Kaiser, and Illia Polosukhin. Attention is all you need. Advances in neural information processing systems, 30, 2017.

[42] Jingjing Xu, Xu Sun, Zhiyuan Zhang, Guangxiang Zhao, and Junyang Lin. Understanding and improving layer normalization. Advances in Neural Information Processing Systems, 32, 2019.

[43] Djork-Arné Clevert, Thomas Unterthiner, and Sepp Hochreiter. Fast and accurate deep network learning by exponential linear units (elus). arXiv preprint 1511.07289, 2015.

[44] Jascha Sohl-Dickstein, Eric A Weiss, Niru Maheswaranathan, and Surya Ganguli. Deep unsupervised learning using nonequilibrium thermodynamics. March 2015.

[45] Alex Nichol and Prafulla Dhariwal. Improved denoising diffusion probabilistic models. February 2021.

[46] Prafulla Dhariwal and Alex Nichol. Diffusion models beat GANs on image synthesis. May 2021.

[47] Robin Rombach, Andreas Blattmann, Dominik Lorenz, Patrick Esser, and Björn Ommer. High-Resolution image synthesis with latent diffusion models. December 2021.

[48] Jonathan Ho and Tim Salimans. Classifier-Free diffusion guidance. July 2022.

[49] Chitwan Saharia, William Chan, Saurabh Saxena, Lala Li, Jay Whang, Emily Denton, Seyed Kamyar Seyed Ghasemipour, Burcu Karagol Ayan, S Sara Mahdavi, Rapha Gontijo Lopes, Tim Salimans, Jonathan Ho, David J Fleet, and Mohammad Norouzi. Photorealistic Text-to-Image diffusion models with deep language understanding. May 2022.

[50] Nan Liu, Shuang Li, Yilun Du, Joshua B Tenenbaum, and Antonio Torralba. Learning to compose visual relations. November 2021.

[51] Nan Liu, Shuang Li, Yilun Du, Antonio Torralba, and Joshua B Tenenbaum. Compositional visual generation with composable diffusion models. June 2022.

[52] Yilun Du, Conor Durkan, Robin Strudel, Joshua B Tenenbaum, Sander Dieleman, Rob Fergus, Jascha Sohl-Dickstein, Arnaud Doucet, and Will Grathwohl. Reduce, reuse, recycle: Compositional generation with Energy-Based diffusion models and MCMC. February 2023.

[53] Yang Song, Jascha Sohl-Dickstein, Diederik P Kingma, Abhishek Kumar, Stefano Ermon, and Ben Poole. Score-Based generative modeling through stochastic differential equations. November 2020.

[54] Berthy T Feng, Jamie Smith, Michael Rubinstein, Huiwen Chang, Katherine L Bouman, and William T Freeman. Score-Based diffusion models as principled priors for inverse imaging. April 2023.

[55] Jiakun Fu, Konstantin F Willeke, Pawel A Pierzchlewicz, Taliah Muhammad, George H Denfield, Fabian Hubert Sinz, and Andreas S Tolias. Heterogeneous orientation tuning across Sub-Regions of receptive fields of V1 neurons in mice. February 2022.

[56] Jia Deng, Wei Dong, Richard Socher, Li-Jia Li, Kai Li, and Li Fei-Fei. ImageNet: A large-scale hierarchical image database. In 2009 IEEE Conference on Computer Vision and Pattern Recognition, pages 248–255, June 2009.

